# Anatomy and Development of the Pectoral Fin Vascular Network in the Zebrafish

**DOI:** 10.1101/2021.04.02.437283

**Authors:** Scott Paulissen, Daniel Castranova, Shlomo Krispin, Margaret Burns, Brant M. Weinstein

## Abstract

The pectoral fins of teleost fish are analogous structures to human forelimbs, and the developmental mechanisms directing their initial growth and patterning are conserved between fish and tetrapods. The forelimb vasculature is critical for limb function, and it appears to play important roles during development by promoting development of other limb structures, but the steps leading to its formation are poorly understood. In this study, we use high-resolution imaging to document the stepwise assembly of the zebrafish pectoral fin vasculature. We show that fin vascular network formation is a stereotyped, choreographed process that begins with the growth of an initial vascular loop around the pectoral fin. This loop connects to the dorsal aorta to initiate pectoral vascular circulation. Pectoral fin vascular development continues with concurrent formation of three elaborate vascular plexuses, one in the distal fin that develops into the fin ray vasculature and two near the base of the fin in association with the developing fin musculature. Our findings detail a complex yet highly choreographed series of steps involved in the development of a complete, functional organ-specific vascular network.

**SUMMARY STATEMENT:** The stereotyped assembly of the pectoral fin vasculature is documented from first migratory sprout into the limb bud, to the adult-like form of the four week old larva.

## INTRODUCTION

The human forelimb is an essential part of the success of the human species. Unfortunately, major defects in forelimb development affect 2.3 out of 10000 live births (Goldfarb et al. 2017). The biological causes of these defects are not all understood, underscoring the importance of studying the multitude of developmental processes that lead to formation of the mature appendage (Goldfarb et al.; Rodríguez-Niedenführ et al., 2001). Fortunately, there are excellent animal models for studying the development of the human arm. Studies on forelimb development in chick (Bates et al., 2002), mouse (Martin, 1990), frog (Abu-Daya et al., 2011), zebrafish (Grandel and Brand, 2011; Isogai et al., 2001; Mercader, 2007), and other vertebrates have shown that the genetic and molecular pathways used to pattern different forelimbs are well conserved. These include evolutionarily conserved gradients of SHH and FGF signaling that provide the molecular basis for orienting the posterior-anterior and proximo-distal axes of the limb (Chiang et al., 2001; Crossley et al., 1996; Mateus et al., 2020; Niswander and Martin, 1992; Nomura et al., 2006; Riddle et al., 1993), as well as important transcription factors like hoxA13, hoxD13 and meis1 involved in regional patterning of the limb (Mercader et al., 2000; Tulenko et al., 2017). When compared, even appendages as divergent as fish pectoral fins and human arms are well conserved at the molecular level (Tulenko et al., 2017).

The development of the forelimb of begins prior to that of the rear limbs and starts as a small tissue protrusion, called the limb bud, on the side of the lateral plate mesoderm (Gros and Tabin, 2014). From this bump, an appendage builds outward beneath the apical ectodermal ridge (AER) (Saunders, 1948). The AER is a thin ridge of tissue that remains on the distal tip of the forelimb that promotes the growth of the limb bud mesenchyme. This mesenchyme becomes the endoskeletal disc (Grandel and Schulte-Merker, 1998). In mammals and fish, the migratory muscle progenitor cells that become the muscles of the forelimb migrate into the limb bud from the ventro-lateral edge of the nearby somites that then differentiate into mature skeletal muscle (Thorsen and Hale, 2005). In chick and mouse, the nerves and blood vessels invade the limb bud on the anterior and posterior edges of the endoskeletal disk and the vessels are thought to be sprouts off of the dorsal aorta (Crossley et al., 1996; Martin, 1990). Although in many respects development of the pectoral fin echoes that of the tetrapod forelimb, one key difference between fin and tetrapod limb development is that the AER elongates extensively in fish to form the large distal portion of the fin supported by cartilaginous fibers known as actinotrichia, whereas in the tetrapod limb bud the AER reduces and condenses into digits (Zhang et al., 2010). As the fin matures, regions of the AER form an array of paired bony, hemi-cylindrical and segmented fin-rays instead of cylindrical segmented digits.

Various aspects of fin development have been studied in the zebrafish, including bone formation (Akiva et al., 2019; Dodo et al., 2020), muscle development (Thorsen and Hale, 2005), neuronal patterning (Dodo et al., 2020; Thorsen and Hale, 2007), regeneration (Marques et al., 2020; McMillan et al., 2013; McMillan et al., 2018; Xu et al., 2014), endothelial cell behavior (Xu et al., 2014) and evolution of the fin-to-limb transition, among others (Lalonde and Akimenko, 2018; Petersen and Ramsay, 2020; Yano et al., 2012; Zhang et al., 2010). Development of the zebrafish vasculature has been studied extensively (Aydogan et al., 2015; Betz et al., 2016; Fujita et al., 2011; Gore et al., 2012; Hasan et al., 2017; Isogai et al., 2001; Lawson and Weinstein, 2002; Reischauer et al., 2016; Torres-Vázquez et al., 2004; Weinstein, 1999; Xu et al., 2014). The study of the vasculature of the fin, however, has mainly in the context of the caudal fin rather than the pectoral fins. Blood vessels associated with the caudal fin-ray bones form stereotypic structure where a single artery flows through the center channel of the paired hemi-cylindrical bones, while two veins drain blood back to the base of the fin ray parallel to the bone in the same plane as the fin itself (Akiva et al., 2019; Huang et al., 2003; Tu and Johnson, 2011; Xu et al., 2014). Other capillary vessels are sparsely found in the membranous region between the bony fin rays, often branching from the artery of one ray and connecting to the vein of an adjacent ray (Huang et al., 2003). The development of the vasculature that feeds the forelimb is comparatively understudied. The initial primitive circulation through the embryonic pectoral fin has been briefly described in previous reports (Grandel and Schulte-Merker, 1998; Isogai et al., 2001; McMillan et al., 2013; Yano et al., 2012). At approximately three days post fertilization (dpf) blood flows from the dorsal aorta to the ventral edge of the pectoral fin bud, the loops around the edge of the pectoral fin until reaches the dorsal edge of the fin bud, where it empties into the common cardinal vein (CCV). The developmental process that leads to the formation of the simple initial embryonic primitive pectoral fin vessel and its extraordinarily extensive elaboration and growth to form the highly complex adult pectoral fin vasculature has not been reported.

The importance of a well-developed, functional forelimb vasculature for proper growth and development of the limb/fin as a whole been highlighted by research on the teratogenic drug thalidomide. It has been known for many years that exposure of embryos to thalidomide during a critical window of limb development causes severe disturbance to or failure of limb formation (Kajii et al., 1973; Vargesson, 2013), and at least a major part of these teratogenic effects appear to be through anti-angiogenic effects of this drug (Therapontos et al., 2009), which has led to exploration of the use of thalidomide and thalidomide analogs as potential anti-angiogenic anticancer agents (Rehman et al., 2011). Zebrafish research has shown a correlation between the teratogenicity of thalidomide analogs and their anti-angiogenic capacity (Mahony et al., 2013), further reinforcing this idea.

In this study, we document the development of the zebrafish pectoral fin vasculature from its first sprouts from the CCV in the zebrafish embryo to an adult-like pectoral fin vascular network in approximately one month-old juvenile fish. Formation of the pectoral fin vasculature involves a highly stereotyped and reproducible stepwise series of events. These include sprouting and growth of an initial vascular loop around the pectoral fin, connection of this loop to arterial blood flow to initiate pectoral vascular circulation, development of an elaborate distal vascular plexus in conjunction with condensation and growth of the bony fin rays, and assembly of two parallel proximal vascular networks near the base of the fin in association with the endoskeletal disk. The successive remodeling and complex elaboration of the pectoral fin vasculature involves a number of interesting events including shunting and re-routing of blood flow at different steps, plexus formation followed by plexus pruning and remodeling, among other fascinating processes. Together, our findings detail a complex yet highly choreographed series of steps involved in the development of a complete, functional organ-and tissue-specific vascular network.

## RESULTS

### Vascular anatomy of the adult zebrafish pectoral fin

The pectoral fin of the adult zebrafish (**Fig. 1A,B**) is a complex structure supplied with oxygen and nutrients by an equally complex vascular network (**Fig. 1C-I**). In terms of its gross anatomy, the pectoral fin can be divided into two major regions-a membranous distal region supported by 11-13 fin rays (lepidotrichia), and a muscular base near the body wall of the zebrafish from which the membranous distal region emerges (**Fig. 1A,B**). The thick proximal portion of the pectoral fin is comprised primary of muscle and bones that articulate and anchor the pectoral fin to the pectoral girdle; its depth and opacity make it difficult to image in the adult. In contrast, the thinner fin-ray containing distal region and its elaborate vascular network are readily visualized using high-resolution optical imaging. Viewed in profile, each fin ray is comprised of two closely apposed hemicylindrical bones that form a hollow tube between them (**Fig. 1C**, (Huang et al., 2003)). Transmitted light imaging of blood circulation in intact adult fins, and tiled confocal imaging of vessels dissected adult fins visualized using the pan-endothelial *Tg(fli1a:egfp)* reporter line (Lawson and Weinstein, 2002), reveal blood vessels running in the space between and closely along either side of the fin ray bones, as well as a sparse network of inter-ray vessels running through the membranous tissues between the fin rays (**Fig. 1D-I** and **Movie 1**). The pectoral fins of male and female zebrafish are largely similar, but a few sex-specific differences are noted. Fin rays (and the vessels within and alongside them) undergo proximal-to-distal branching, and while the terminal distal fin ray bifurcations occur at approximately the same distance from the distal end of the pectoral fin in males and females, the penultimate branches of the fin rays and fin ray vessels occur at a more proximal position in male pectoral fins (**Fig. S1**, white arrows). Male pectoral fins (**Fig. 1D-F** and **Fig. S1**, cyan arrows) also have a higher density of inter-ray vessels than female pectoral fins (**Fig. 1G-I** and **Fig. S1**, magenta arrows). As has been previously reported (McMillan et al., 2013), male pectoral fins also have a dense network of vessels associated with the breeding tubercles, specialized toothlike keratinized epidermal protrusive structures clustered on the dorsal surface of male pectoral fins that are thought to be involved in helping maintain contact during spawning (**Fig. S1**)(McMillan et al., 2015).

**Figure 1.**
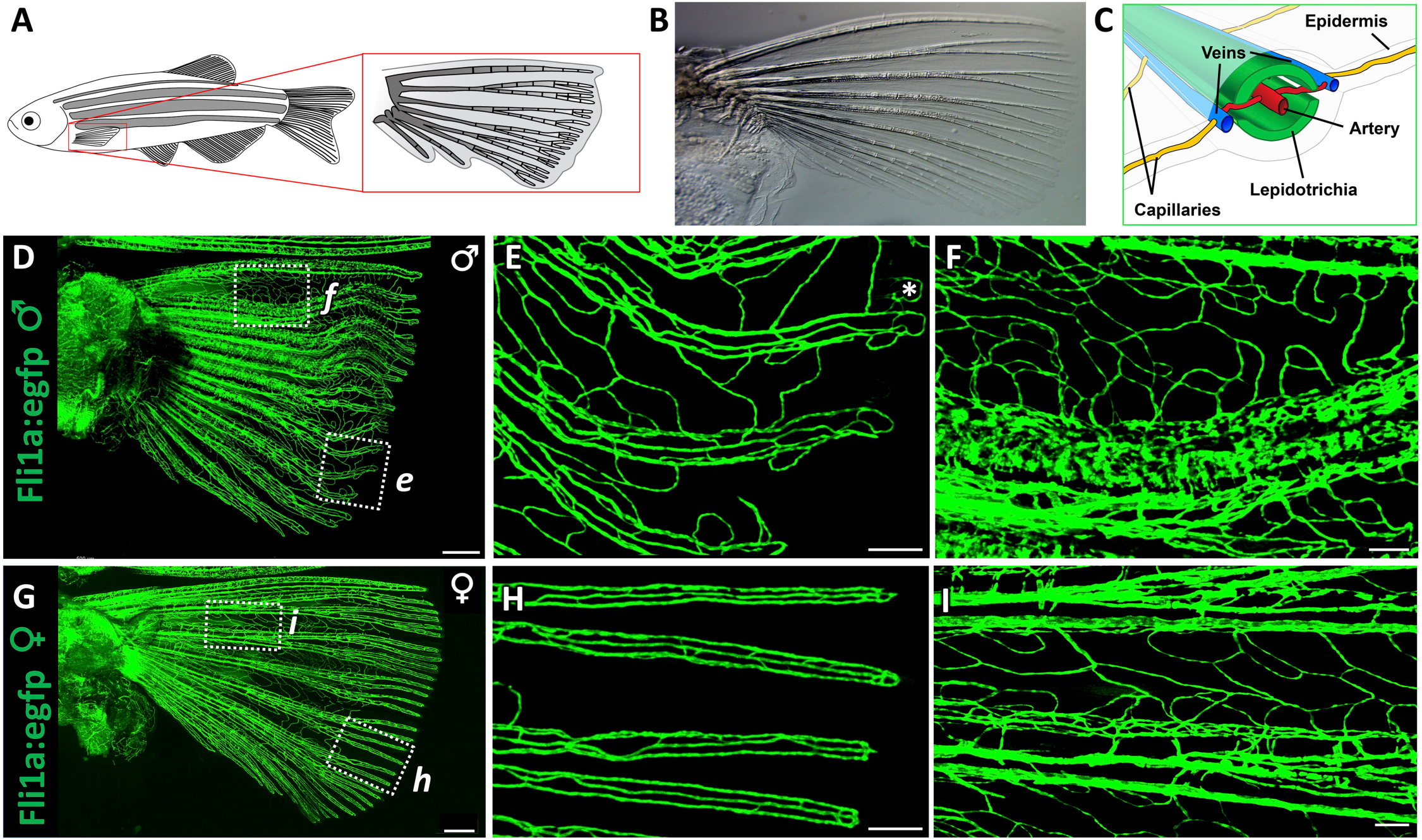
Vascular anatomy of the adult zebrafish pectoral fin. (**A**) Drawing of an adult zebrafish with a magnified view of the pectoral fin and its lepidotrichia (fin ray bones). (**B**) Transmitted light image of a dissected pectoral fin. (**C**) Schematic diagram showing a cross-section of a zebrafish fin ray. (**D-I**) Tiled confocal images of pectoral fins dissected from adult male (D-F) or adult female (G-I) *Tg(fli1a:egfp)* transgenic zebrafish. Panels E and F show higher magnification views of the boxed regions in panel D. Panels H and I show higher magnification views of the boxed regions in panel G. A, Fin ray artery; V, paired fin ray vein; I, fin ray interlinking vessels. Scale bars = 500 μm (B,D,G), 150 µm (E,H,), 100 µm (F,I).

The vessels inside the fin rays generally flow in a proximal-to-distal direction, while the paired vessels along either side of the fin rays generally flow in a distal-to-proximal direction, suggesting that these represent arterial and venous vessels, respectively (**Fig. 2A** and **Movie 1**). The arterial identity of the vessel running down the center of each fin ray is supported by confocal imaging of the fin vasculature using the arterial *Tg(kdrl:mcherry)* transgenic line. In medial areas of the fin, mCherry-positive arterial vessels can be seen running down the center of each weakly blue autofluorescent fin ray (**Fig. 2B-E**), while the lateral vessels juxtaposed to the fin rays and inter-ray vessels (see **Fig. 1**) are only weakly mCherry-positive or mCherry-negative. In more distal areas of the fin the fin ray arteries are only weakly mCherry-positive, although interestingly small lateral segments projecting from these distal arteries remain more strongly mCherry-positive (**Fig. 2B,F-I**).

**Figure 2.**
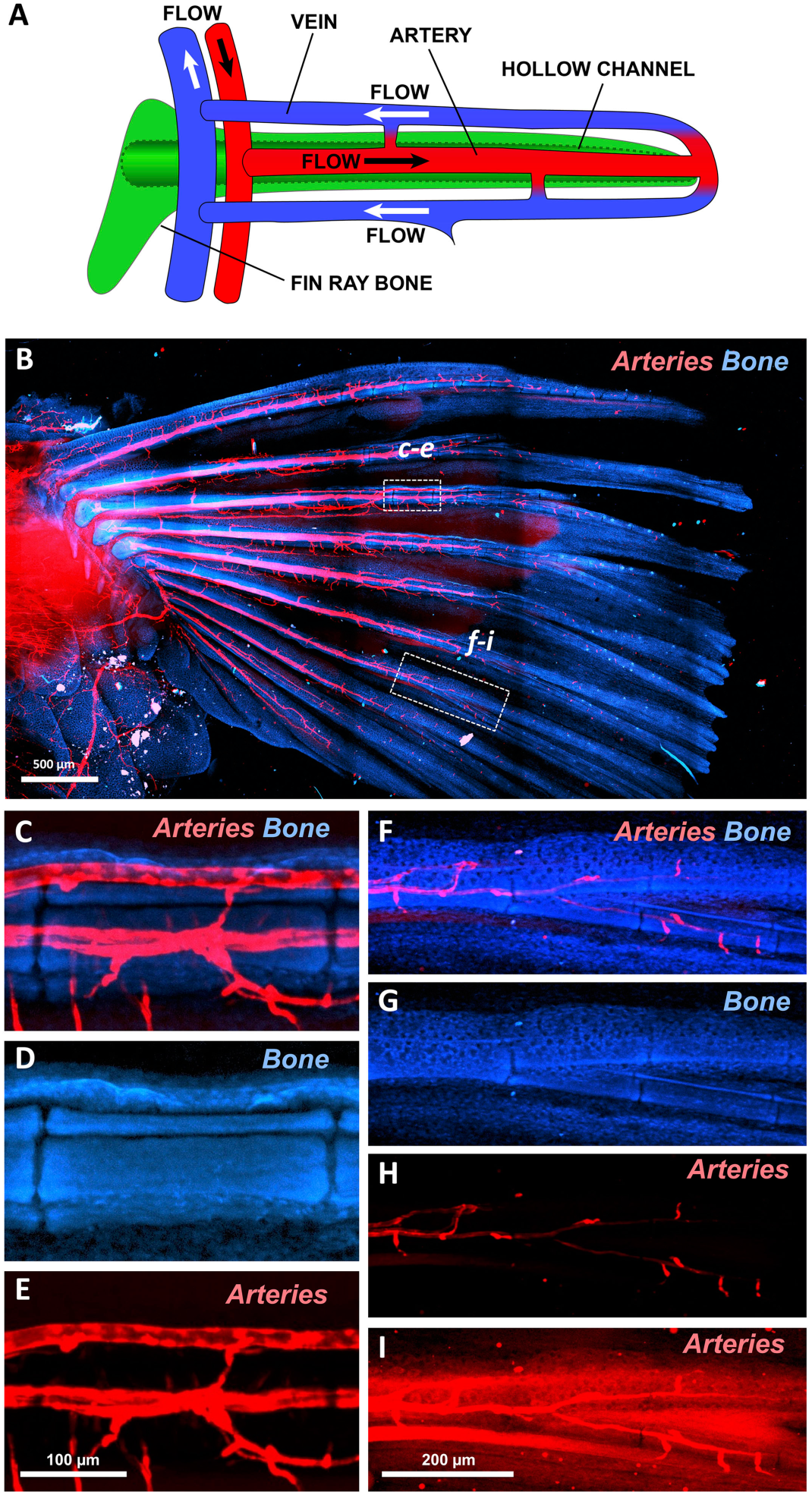
Fin ray-associated vasculature of the adult zebrafish pectoral fin. (**A**) Schematic diagram illustrating circulatory flow through an individual adult zebrafish fin ray. Arteries are in red, veins in blue, and fin ray bone in green. (**B-I**) Confocal images of an adult pectoral fin dissected from a *Tg(kdrl:mcherry)* zebrafish, showing kdrl:mcherry-positive arterial blood vessels (red) and autofluorescent bone (blue). (B) Low-magnification tiled confocal image of a dissected adult pectoral fin. (C-E) Higher-magnification confocal images of boxed region *(c-e)* in panel B, showing kdrl:mcherry-positive arteries (C,E) running through the center of autofluorescent bones (C,D) in the proximal region of a fin ray. (F-I) Higher-magnification confocal images of boxed region *(f-i)* in panel B, showing weakly kdrl:mcherry-positive arteries (F,H,I) running through the center of autofluorescent bones (F,G) in the distal region of a fin ray. Panel I shows an “overexposed” image of the fluorescence in Panel H. (**J**) Simplified schematic diagram illustrating some of the major circulatory pathways in the adult zebrafish pectoral fin. Scale bars = 500 μm (B), 150 µm (C-E), 200 µm (F-I).

The fin ray central arteries and paired lateral veins are connected to arterial feed and venous drainage at the base of each fin ray (**Fig. 2A**). The arterial feed for the fin vasculature is supplied by a vascular branch emerging from the dorsal aorta, while venous drainage is accomplished primarily via the common cardinal vein (described further below). In addition to the more distal fin ray vasculature, the adult fin also contains an elaborate plexus of vessels in the proximal area of the fin below the base of the fin rays (**Fig. 1D,G**, arrows). As also described below, this proximal vascular plexus provides at least a portion of the arterial feed for the distal fin ray arteries (**Fig. S1**, yellow arrows). The vasculature of the pectoral fin becomes more complex near the base of the fin rays, and particularly so in the proximal plexus below the fin rays, although as noted above imaging of the proximal regions of the adult pectoral fin is extremely challenging due to the thickness and opacity of this bony and muscular tissue. In the sections below we describe the stepwise development of the distal and proximal vascular networks of the pectoral fin from the first vascular sprouts in the fin bud at approximately 1.5 day old embryo to a complex vascular network with most of the major features of the adult pectoral fin vascular network in the 4-week-old juvenile zebrafish

### Pectoral fin vascular development: Formation and perfusion of the primitive pectoral artery

Initial assembly of the vascular network of the pectoral fin follows a highly stereotyped plan (**Fig. 3**, and **Movie 2**). Pectoral fin vascular development begins at approximately 30-34 hours post-fertilization (hpf) with the emergence of a pair of dorsal and ventral vascular sprouts from the caudal face of the common cardinal vein (CCV)(**Fig. 3A,D** and **Movie 2**). These sprouts invade and then elongate along the rim of the growing fin bud (**Fig. 3B,E** and **Movie 2**), eventually joining together at the posterior end of the fin to form a complete primitive pectoral artery by approximately 38-40 hpf (**Fig. 3C,F** and **Movie 2**). The primitive vascular arc that becomes the pectoral artery initially possesses only venous connections to the CCV at either end and does not immediately lumenize or carry blood flow.

**Figure 3.**
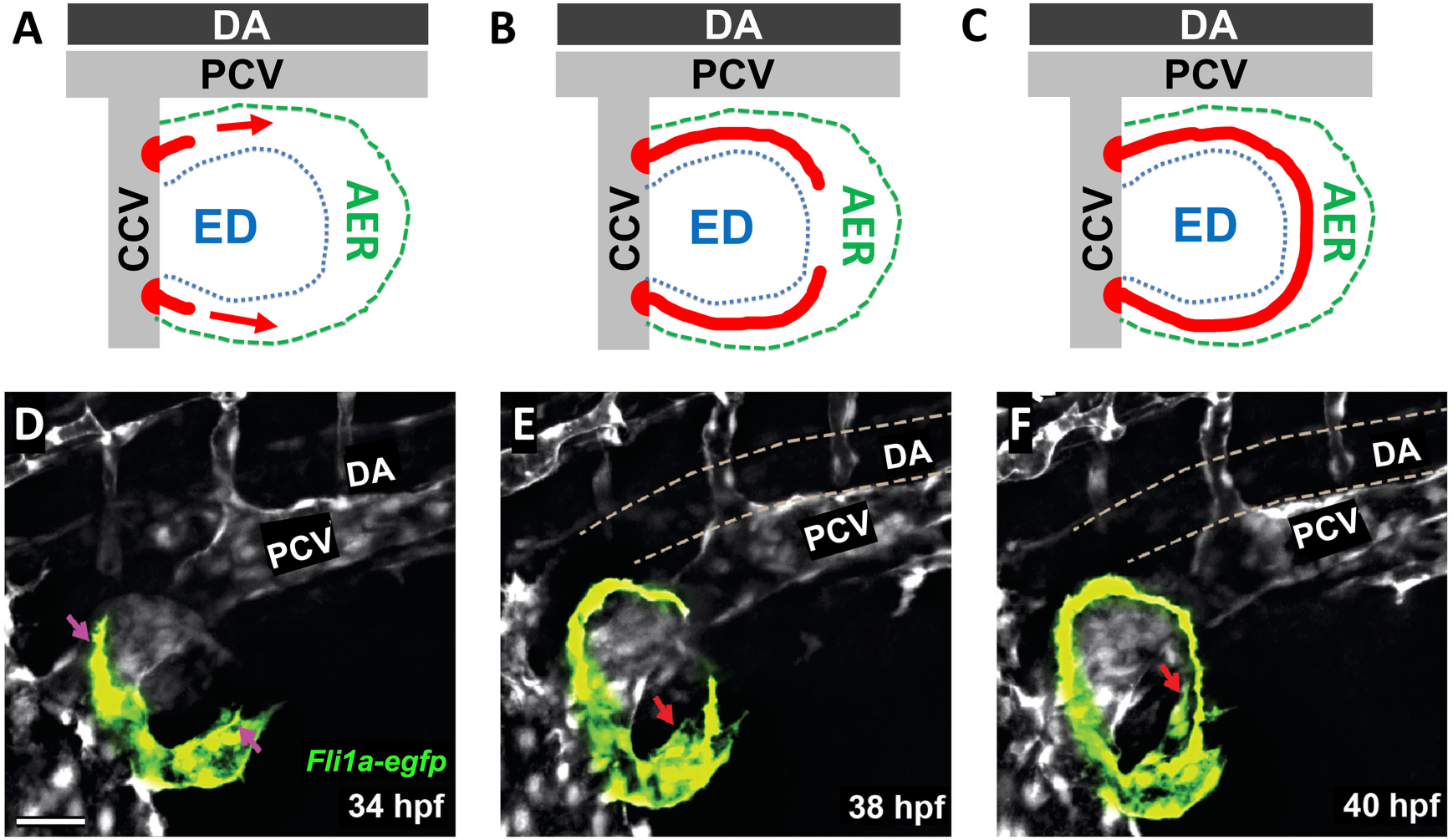
Formation of the primitive pectoral artery. (**A-C**) Schematic diagrams illustrating stages of primitive pectoral artery formation, including initial dorsal and ventral sprouts from the CCV (A), elongation of these sprouts along the rim of the growing fin (B), and linkage of the two segments to form a complete primitive pectoral artery vascular arc (C). (**D-F**) Still images taken from a time lapse confocal time series of primitive pectoral artery formation in a *Tg(kdrl:egfp)* transgenic zebrafish at 34 hpf (D), 38 hpf (E), and 40 hpf (F). See **Movie 2** for complete time-lapse series corresponding to panels D-F. Scale bars = 40 μm (D-F).

Shortly after or concomitantly with completion of the primitive pectoral fin artery vascular arc, a new vascular sprout emerges from the pectoral artery close to its ventral connection point with the CCV (**Fig. 4A,D**, and **Movie 2**). This sprout grows medially, dorsally, and caudally (**Fig. 4B,E**, and **Movie 2**), traveling approximately ∼300 microns in total and migrating underneath and between the somites before eventually connecting to the dorsal aorta at the base of the second intersegmental vessel, or to the second ISV itself (the second ISV is always arterial – see Isogai et al., 2001)**(Fig. 4C,F)**. Once this connection is made flow begins (**Fig. 4G-U**, and **Movie 2**). In most cases a ventral connection directly to the CCV is still present when flow initiates, causing blood to shunt directly into the CCV through this ventral connection, leaving the majority of the primary arc unperfused (**Fig. 4H,N-Q**, and **Movies 3,4,5**). However, this ventral connection is generally severed within a few hours and flow is redirected through the pectoral artery primary arc, emptying into the CCV via the dorsal connection only (**Fig. 4I,R-U**, and **Movies 3,4,5**). The initial ventral shunt to the CCV is present in at least 63.6% of fish observed (*n*=11), persisting 110 ± 95 min before being severed (**Fig. S2**). The remainder of the time either the entire primary arc perfuses immediately without a shunt to the CCV, or the shunt disconnects too rapidly to be observed in our time-lapse imaging. The average onset of flow through the complete pectoral artery arc occurs at 63 ± 5 hpf and the endothelial cells bridging the former ventral connection to the CCV is pruned, preventing further resumption of shunting (**Fig. S2** and **Movies 3,5**).

**Figure 4.**
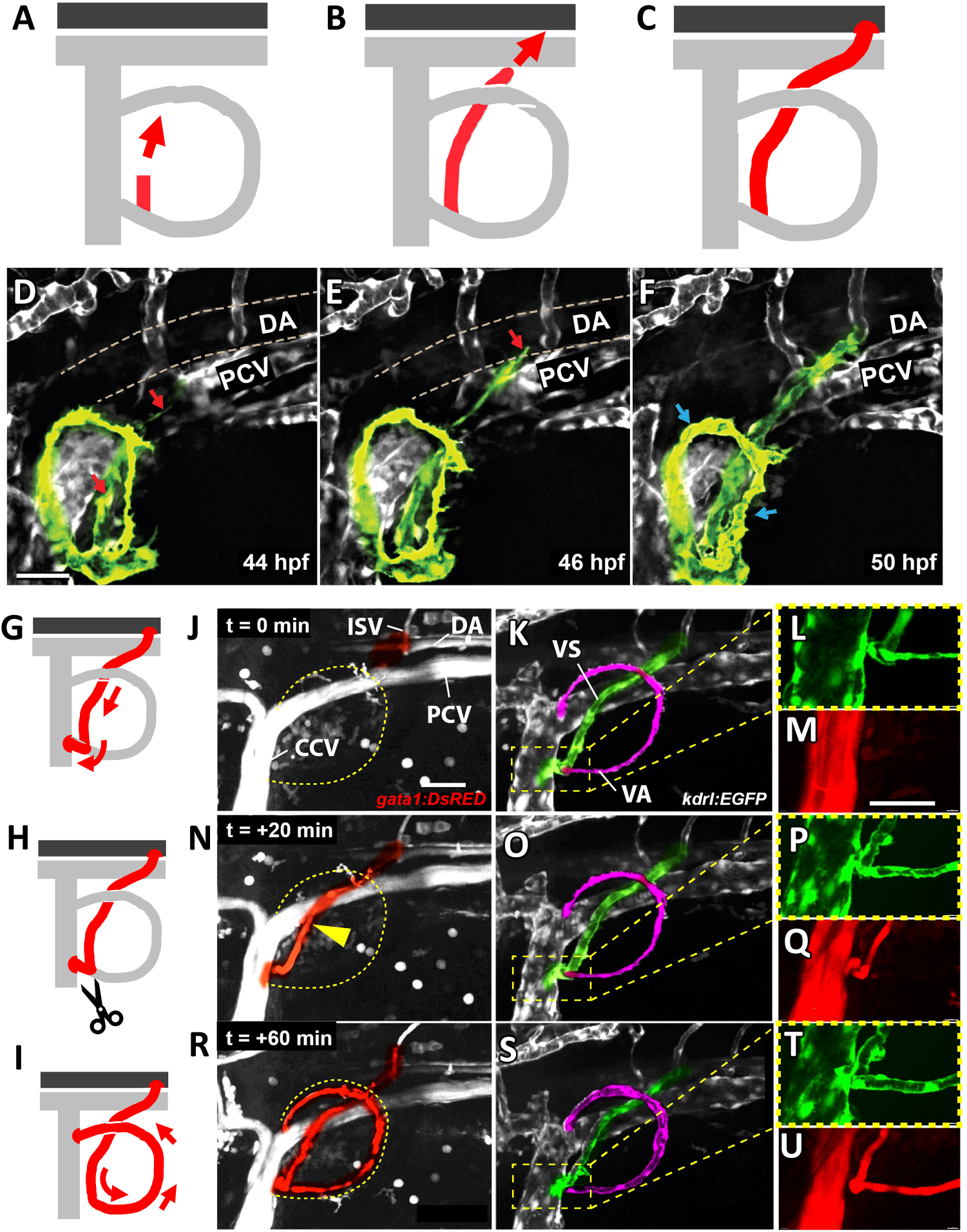
Perfusion of the primitive pectoral artery. (**A-C**) Schematic diagrams illustrating stages in the formation of a connecting link between the primitive pectoral artery and the dorsal aorta, including formation of a new sprout from the base of the primitive pectoral artery where it connects ventrally to the CCV (A), medial and dorsal-caudal elongation of the new vessel segment (B), and linkage to the dorsal aorta at or near the base of the second intersegmental vessel (C). (**D-F**) Confocal still images from a time-lapse series showing formation of a connecting link between the primitive pectoral artery and the dorsal aorta in a *Tg(kdrl:egfp)* transgenic zebrafish at 44 hpf (D), 46 hpf (E), and 50 hpf (F). See **Movie 2** for the complete time-lapse series corresponding to panels D-F. (**G-I**) Schematic diagrams illustrating initial shunting of flow from the DA through the base of the pectoral fin directly to the CCV (G), followed by disconnection of this shunt (H) and re-routing of flow through the primitive pectoral artery primary arc (I). (**J-R**) Confocal images from a time-lapse series of the pectoral fin and adjacent structures in a 2-day old *Tg(gata1:dsred), Tg(kdrl:egfp)* double transgenic zebrafish, showing gata1:dsred-positive blood cells (J,L,M,O,P.R) and kdrl:egfp-positive blood vessels (K,L,N,O,Q,R). Panels L,O,R show higher magnification gata1:dsred (red)/kdrl:egfp (green) confocal images of the boxed regions in panels K,N,Q, respectively. (**J-L, t=0 min**) The vascular link to the dorsal aorta has assembled (highlighted green in panel K) but has not yet lumenized, and no blood flow is present. (**M-O, t=20 min**) The connection to the dorsal aorta (highlighted green in panel N) has been completed and blood is now flowing through this connecting vessel (highlighted red in panel M), but the blood is being shunted directly to the CCV without perfusing the primitive pectoral artery arc (highlighted magenta in panel N). (**P-R, t=40 min**) The shunt to the CCV has now disconnected and all blood flow has been rerouted into the primitive pectoral artery arc (highlighted red in panel P). Blood is only emptying into the CCCV through the dorsal connection. See **Movie 3** for the complete time-lapse series corresponding to panels J-U. See **Movies 4** and **5** for higher magnification images and 3-D reconstructions of the ventral junction to the CCV and its shunting and then disconnection. Scale bars = 30 μm (D-F), 40 μm (J-U).

### Pectoral fin vascular development: Formation of the distal fin ray vascular network

Once established, the initial pectoral artery circulatory loop changes little over the next several days, with the exception of a variable degree of sprouting, branching, and formation of a small plexus at the distal tip of the growing fin (**Fig. 5A-F, Movie 8** and **Fig. S3**). Branching and complexity of the distal fin ray vascular plexus begins to increase dramatically, however, at 13.7±1.0dpf, or 5.3±0.3mm in total animal length (**Fig. 5G-K, Movie 8** and **Fig. S3, S4**; note that at later stages fin vessel development correlates less well with chronological age than with animal size, which can vary substantially for animals of the same age (Fig S4)). From approximately ∼7.0 mm (17 dpf) through roughly 9.0 mm (22 dpf), a progressive dorsal-to-ventral remodeling of the distal plexus takes place that leads to the formation of an adult-like fin ray vascular network with well-defined arterial and venous vessels (**Fig. 5L-Q**, and **Movie 8**). As visualized using a *Tg(kdrl:mCherry), Tg(mrc1a:egfp)* double transgenic line, this remodeling begins with localized hypersprouting/hyperbranching in the dorsal-most region of the distal plexus that then spreads to more ventral areas of the plexus at size 7.6±0.3mm or 21.1±2.0 dpf age (**Fig. 5N-Q,, Movie 8** and **S4**). At 7.8mm±0.4mm size or 21.6±2.0 dpf age, a small part of this dorsal hyperbranched plexus begins to develop into an arterial network with persistently kdrl-positive and progressively mrc1a-negative (presumptive arterial) vessels that extend into the developing fin rays. As this occurs, most of the remaining mrc1a-positive plexus regresses, particularly just dorsal to each fin ray, and a single mrc1a-positive, kdrl-negative (presumptive venous) vessel coalesces along the ventral side of each fin ray (**Fig. 5N-R** and **Movie 8**). These initial arterial and venous vessels appear at 8.41mm±0.3mm size or 23.4±2.2 dpf age, and continue to extend along each fin ray as it grows in length (**Fig. 5L,N-Q, S3, S4** and **Movie 8**). In addition, starting at approximately 9.0mm new venous sprouts emerge dorsally from near the end of each developing fin ray and then grow proximally and distally along the dorsal side of the fin rays to give rise to the paired dorsal fin ray veins (**Fig. 5N-Q**, and **Movie 8**). Completion of the process described above results in formation of a very adult-like independent fin ray vasculature with a single arterial feed running down each fin ray and paired venous drains on either side (**Fig. 5P,Q, S3**; see **Fig. 1** and **Fig. 2** for comparison to adult fin ray vessels).

**Figure 5.**
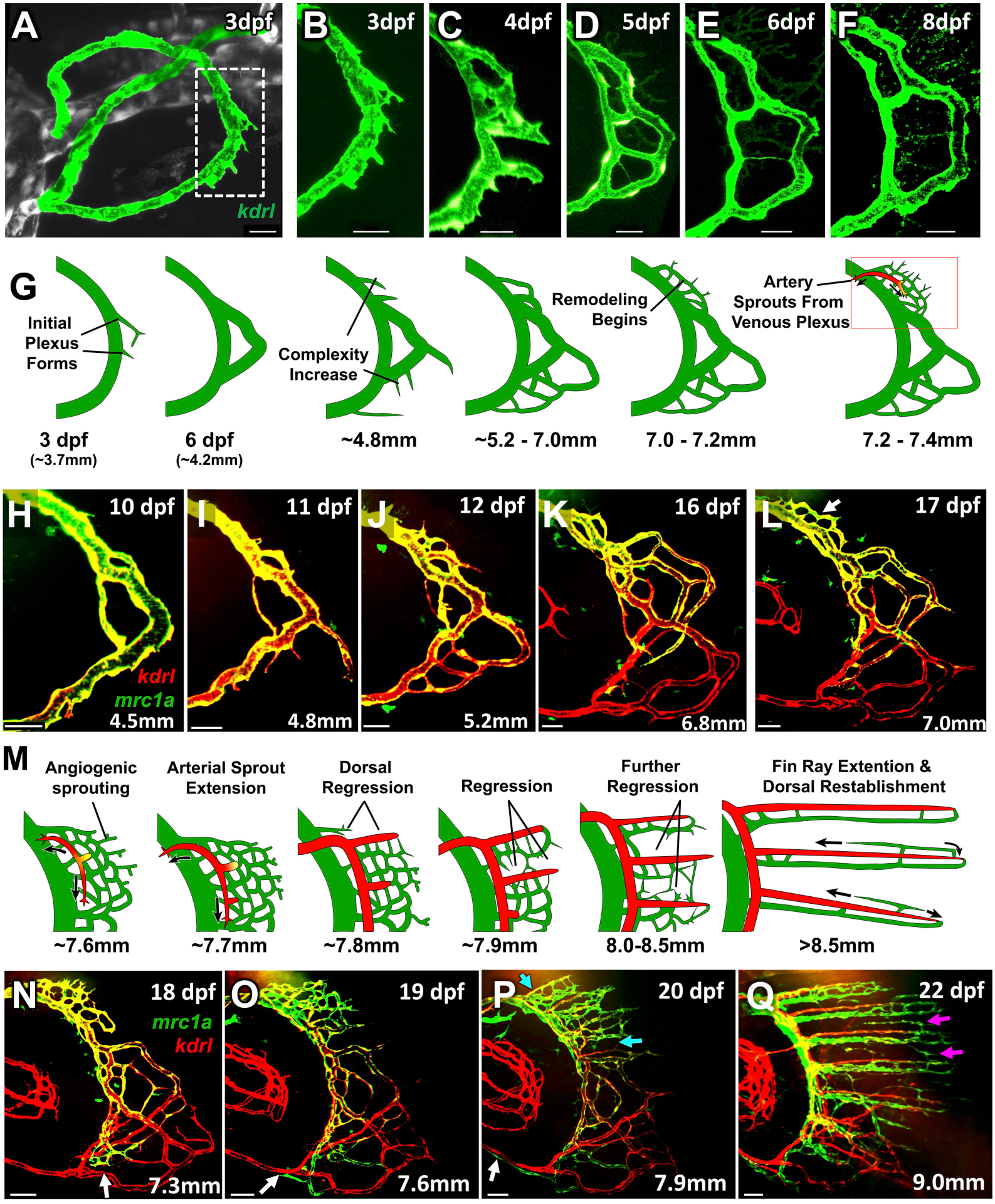
Formation of the distal fin ray vasculature. (**A**) Confocal image of the pectoral fin vessels (green) of a 3 dpf *Tg(kdrl:egfp)* transgenic zebrafish (other surrounding vessels in grey). Dashed box indicates the approximate area imaged in panels B-F. (**B-F)** Confocal image time series of the pectoral fin distal vascular plexus of the same *Tg(kdrl:egfp)* transgenic zebrafish as in panel A, imaged at 3 (B), 4 (C), 5 (D), 6 (E), and 8 (F) dpf. (**G**) Schematic diagram illustrating the series of events occurring in the distal fin from approximately 3 dpf to 18 dpf (up to approximately 7.4 mm), encompassing the stages shown in panels A-L. (**H-L**) Confocal image time series of the pectoral fin distal vascular plexus of the same *Tg(kdrl:mCherry), Tg(mrc1a:egfp)* double transgenic larva imaged at 10 (H), 11 (I), 12 (J), 16 (K), and 17 (L) dpf (4.5, 4.8, 5.2, 6.8, and 7.0 mm in length, respectively). See **Movie 6** for 3-D reconstructions of the image in panel L. (**M**) Schematic diagram illustrating the series of events occurring in the distal fin from approximately 18 dpf to 22 dpf (7.3 to 9.0 mm), encompassing the stages shown in panels N-Q. (**N-Q**) Confocal image time series of the pectoral fin distal vascular plexus of the same *Tg(kdrl:mCherry), Tg(mrc1a:egfp)* double transgenic larva imaged at 18 (N), 19 (O), 20 (P), and 22 (Q) dpf (7.3, 7.6, 7.9, and 9.0 mm in length, respectively). See **Movie 7** for 3-D reconstructions of the images in panels N-Q. A “time series” of the images in panels H-Q is shown in **Movie 8**. All scale bars = 50 μm.

Endothelial hypersprouting, distal plexus remodeling, and formation of the fin ray arteries are temporally and spatially correlated with fin ray bone formation, as visualized in a *Tg(kdrl:egfp), Tg(sp7:ntr-2a-mcherry)* double transgenic line with EGFP and mCherry expression in vessels and bone, respectively (**Fig. 6**). Initial mCherry-positive bone condensations are closely associated with local formation of a hypersprouting, highly branched plexus of smaller vessel segments (**Fig. 5O,P** and **Fig. 6A,B**). As the second fin ray bone begins to coalesce, a new bidirectional angiogenic sprout begins to migrate out from the distal plexus near the base of the future second fin ray bone (**Fig. 6A-C**). The proximal/dorsal tip of this sprout grows into the pectoral fin, eventually connecting with a separate more proximal plexus of vessels beginning to develop in the center of the fin in association with the endoskeletal disk (**Fig. 6C-F**, yellow arrows; formation of the proximal plexus is discussed below). The distal/ventral tip of the sprout grows ventrally, parallel and in close proximity to the original primary pectoral artery, to which it connects near the ventral side of the fin (**Fig. 6C-F**, blue arrows). Branches of this newly developing arterial network grow into each of the fin ray bones in dorsal-to-ventral sequence as the bones develop and extend (**Fig. 6E,F**, asterisks). The developing arterial network maintains persistently strong *kdrl:egfp* expression as the remainder of the distal fin vascular network either regresses (see **Fig. 5P,Q**) or progressively loses *kdrl:egfp* expression (see **Fig. 6E,F, S3**) and remodels into veins (**Fig. 5P,Q**). Once the newly developing arterial network connects to the proximal plexus dorsally and the primitive pectoral artery arc ventrally (yellow and blue arrows in **Fig. 6F**) it begins to receive arterial blood flow from both ends, acting as a “manifold” to supply blood to the fin ray arteries that, in turn, drain into the fin ray veins. The fin ray veins then flow back to another, separate venous manifold developing at the base of the fin rays, that drains dorsally into the CCV. This new venous manifold emerges at least in part from the original dorsal part of the primitive primary pectoral artery arc, extending further ventrally by angiogenic sprouting and growth alongside the developing arterial network (**Fig. 6F**).

**Figure 6.**
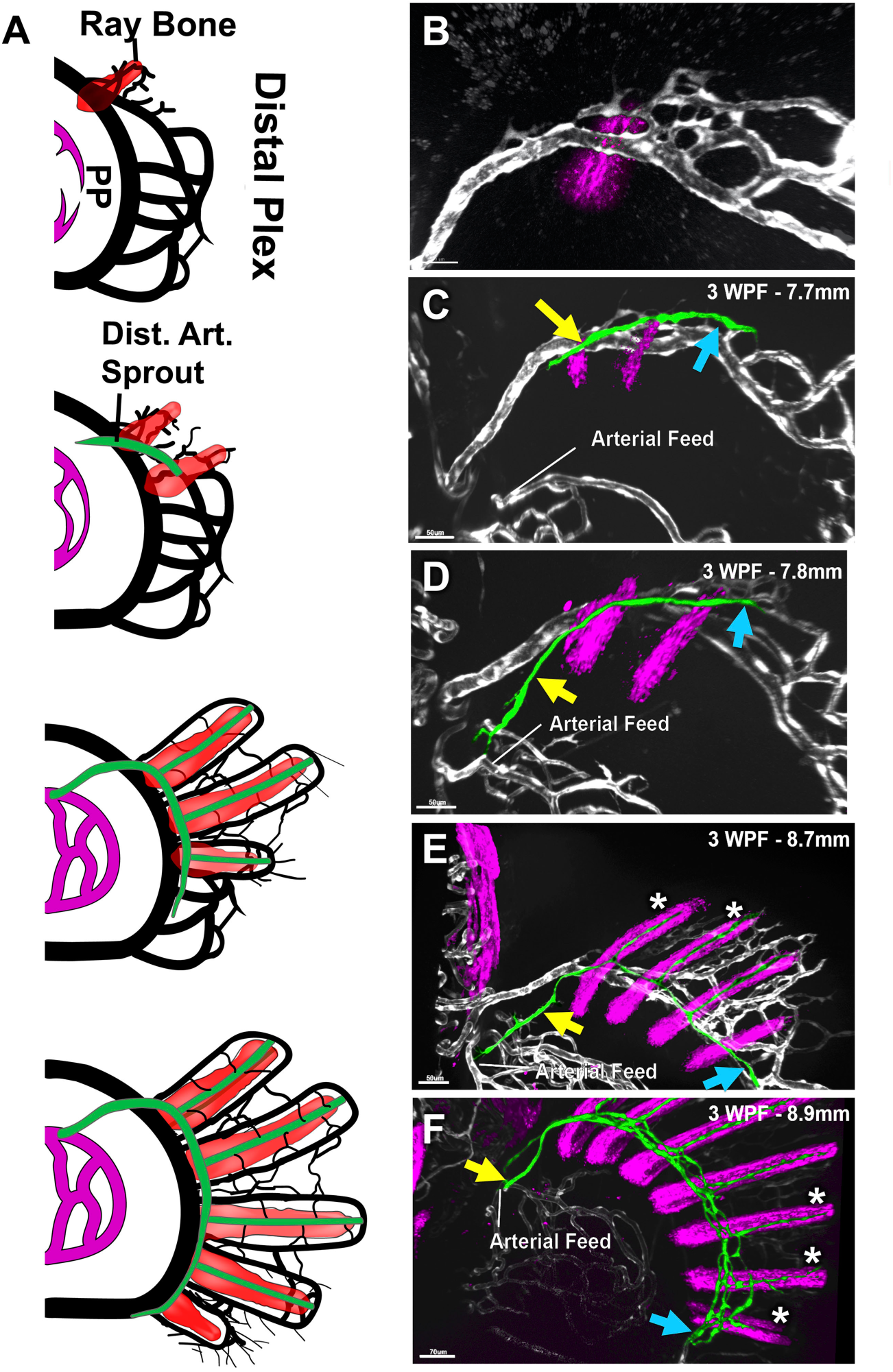
The formation of the distal fin ray arterial network begins as a sprout from the distal venous plexus. (**A**) Schematic illustrating the relationship between the developing fin ray bones (light red) and developing fin ray arterial network (green) that emerges from the pectoral fin distal vascular plexus (black) in approximately 7.0 mm to 9.0 mm fish. (**B-F**) Confocal images of the pectoral fin distal vascular plexuses of separate 3 week old *Tg(sp7:mcherry-ntr), Tg(kdrl:egfp)* double transgenic zebrafish of different sizes, with mCherry-positive condensing bone in magenta and EGFP-positive vessels in grey, except for the emerging fin ray arterial network which is false-colored in green. Larvae shown are 7.4 mm, (B), 7.7 mm (C), 7.8 mm (D), 8.7 mm (E), and 8.9 mm (F) in length. Yellow and blue arrows indicate the proximal/dorsal and distal/ventral ends, respectively, of the developing fin ray arterial vascular network. White asterisks indicate developing individual fin ray arteries. Scale bars = 100 μm.

### Pectoral fin vascular development: formation of the proximal plexus

As development of fin ray vascular network is taking place in the distal regions of the fin, a separate but equally elaborate vascular network is beginning to develop in the proximal part of the fin in association with the endoskeletal disk (**Fig. 7**). At 4.8±0.3mm, sprouts emerge from the pectoral fin artery, the common cardinal vein, and an unidentified venous source rostral to the pectoral fin that invade the lateral and medial sides of the endoskeletal disk (**Fig. 7A-D,S4**, and **Movie 9**). Over the next several days these sprouts rapidly elaborate into two new complex and stratified vascular plexuses (**Fig. 7A,B,E-G**, and **Movie 9**). The two plexuses form within the abductor and adductor muscles on the lateral and medial sides of the endoskeletal disc. Although the detailed vascular pattern of each of these new plexuses is highly variable, they have a number of consistent features. The arterial feed for the medial proximal plexus comes from the same ventral connection to the dorsal aorta that supplies the distal fin ray vascular plexus (**Fig. 7B,G,H,I**), while drainage from the medial plexus is achieved via connections to the common cardinal vein (CCV) (**Fig. 7B,G,I**). The lateral proximal plexus, in contrast, forms drains to a vessel that runs laterally to the CCV and empties into an undetermined high-caliber vessel rostral to the pectoral fin (**Fig. 7H,I**). By 20 dpf the proximal plexus exists as two elaborate vascular plexuses arranged in parallel within the plane of the fin endoskeletal disk, with a limited number of connections between them (**Fig. 5Q**, and **Movie 9**).

**Figure 7.**
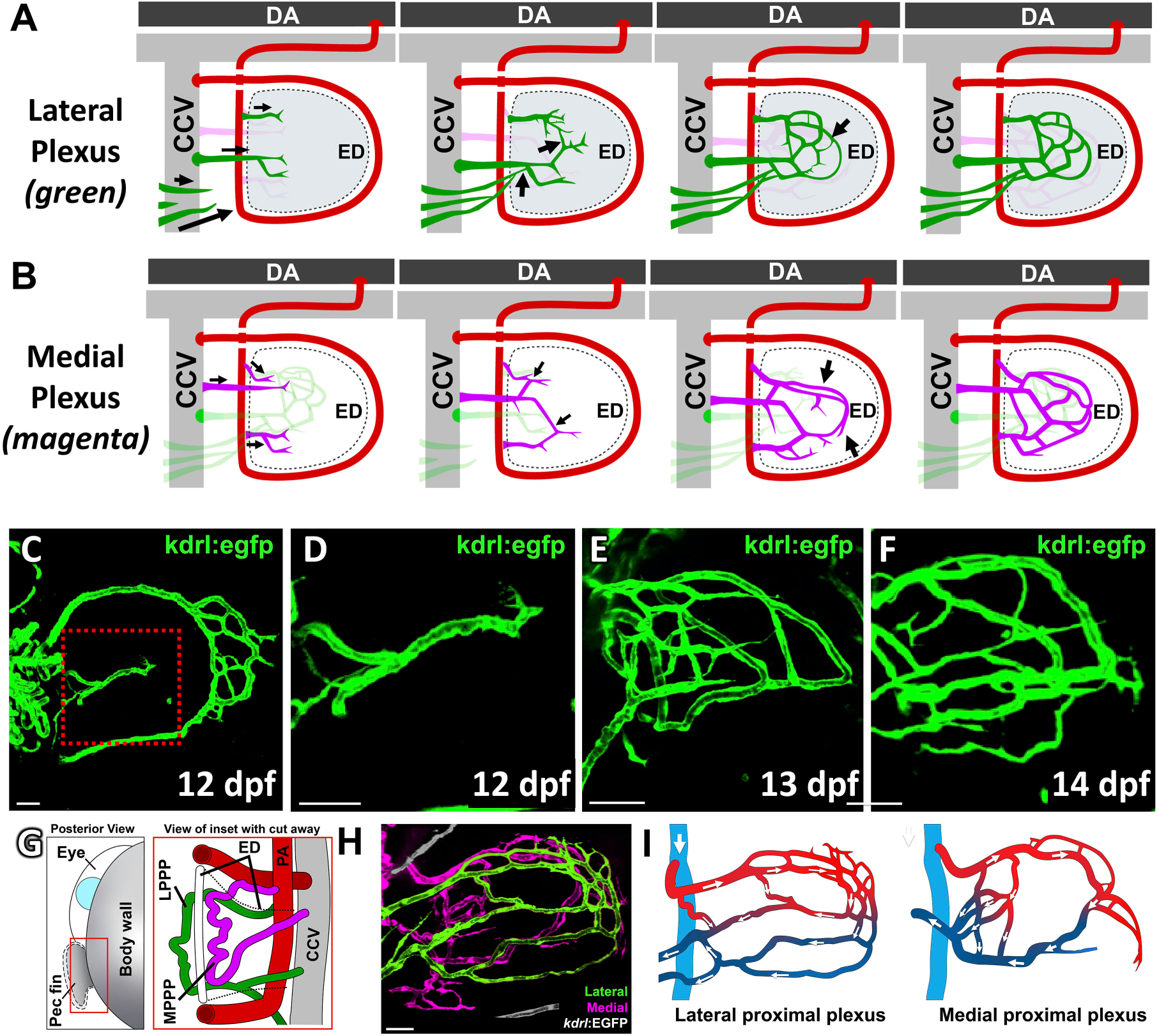
Formation of the proximal vascular plexus. (**A,B**) Schematic diagrams illustrating the growth and development of the lateral (A; in green) and medial (B; in magenta) proximal plexuses in association with the pectoral fin endoskeletal disc, occurring approximately between 6 mm and 8 mm larval length. (**C-F**) Confocal images of the pectoral fin vasculature of three separate *Tg(kdrl:egfp)* larvae at 12 dpf (C,D), 13 dpf, (E), and 14 dpf (F). The red box in panel C indicates the approximate location of the higher magnification images in panels D-F, which show the proximal plexus in progressively older animals. (**G**) Confocal image of an 18 dpf *Tg(kdrl:egfp)* fish with the medial and lateral proximal plexuses false colored in magenta and green respectively. Note the vessels in the lateral plexus that disappear rostrally from the image. See **Movie 9** for 3-D image reconstructions of the same confocal stack shown in the image in panel G. (**H**) Schematic diagrams illustrating the morphology and circulatory flow patterns of the medial (left) and lateral (right) proximal vascular plexuses shown in the confocal image in panel G. Red and blue labeling facilitates visualization of vessels supplying or draining the plexuses. (**I**) Schematic diagram illustrating the positioning of the lateral and medial pectoral proximal plexuses, (LPPP and MPPP, respectively) on either side of the endoskeletal disc within the larval pectoral fin. Scale bars = 50 μm.

Although the vasculature of both the proximal vascular plexus and the fin rays increases further in complexity and the vessels form additional interconnections between the first 3 weeks of development and the adult pectoral fin, the basic arrangement of circulatory flow patterns in the fin is largely maintained into adulthood, although it becomes highly elaborated to accommodate the increased amount of tissue in the adult. However, we take note below of an additional later-stage change to the vascular pattern of the pectoral fin - formation of links between the proximal and distal pectoral fin vasculature.

### Formation of links between the proximal and distal pectoral fin vasculature

Observation of older larval/early juvenile fish reveals later-stage addition of links between the newly developed medial proximal plexus and fin ray arterial networks described above (**Fig. 8A-G**). These links begin to form when the larva is approximately 9.0 mm long, and they extend from the medial proximal plexus to the artery at the base of each fin ray. These links robustly express the *kdrl:mcherry* transgene but express little or no *mrc1a:egfp* and begin to appear first on the dorsal-most portion of the pectoral fin. They form from sprouts that emerge from both the medial proximal plexus and from the artery at the base of the fin rays. Once the inter-plexus links have formed, they become new arterial feeds for the fin ray “arterial manifold;” no connections are observed being made to the “venous manifold” at the base of the fin rays, consistent with the *kdrl:mcherry* transgene-positive-only expression of the new inter-plexus links.

**Figure 8.**
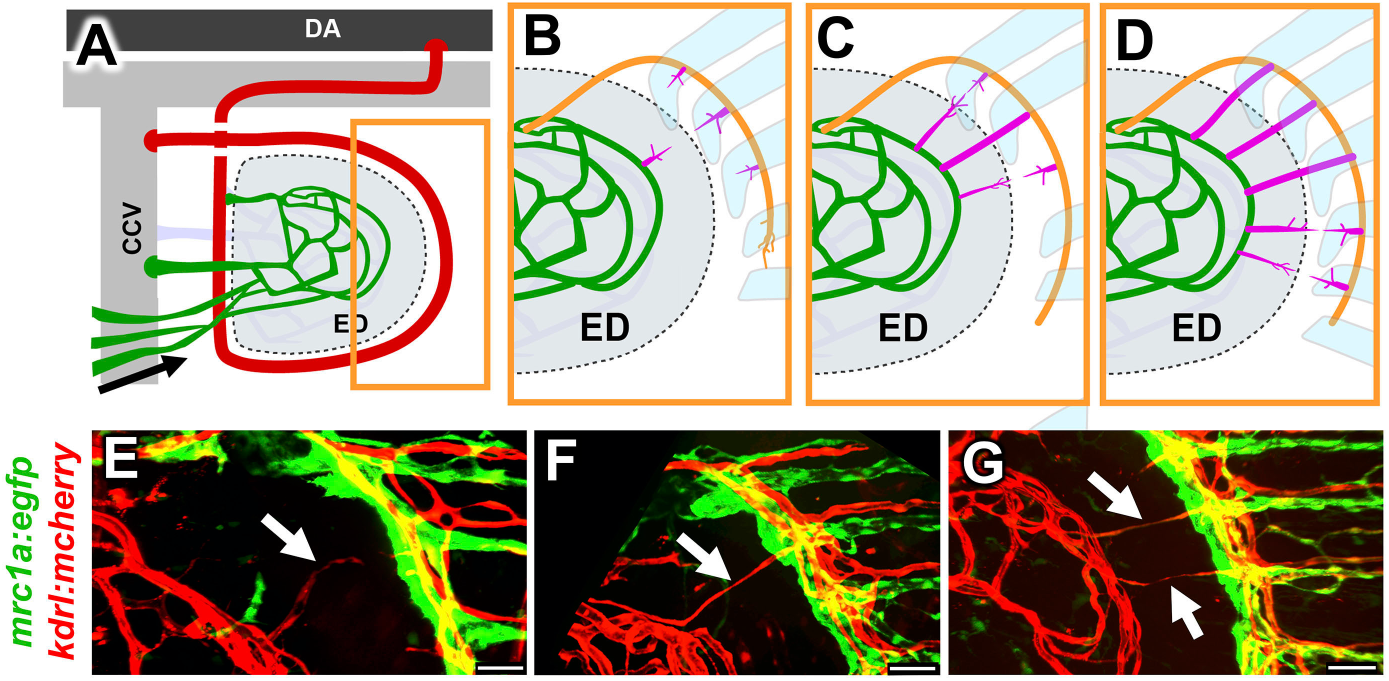
Late-stage remodeling leading to the formation of the interplexus bridges and regression of the dorsal arm of the primary arc. (**A-D**) Schematic diagrams illustrating formation of links between proximal and distal vascular plexuses. (A) Simplified overview diagram of the pectoral fin vasculature at approximately 3.5 weeks, showing the proximal plexus (green) and arterial feed for the distal fin ray vascular network (red). The box shows the approximate magnified area depicted in panels B-D. (B-D) Stages in formation of inter-plexus links. (**E-G**) Confocal images of the inter-plexus region of the pectoral fin of three separate 3.5 week old *Tg(kdrl:mCherry), Tg(mrc1a:egfp)* double transgenic zebrafish, showing the formation of arterial (mCherry-positive, EGFP-negative) links (white arrows) between the medial proximal plexus (to the left) and the arterial fin ray vascular plexus (to the right). Scale bars = 50 μm.

## DISCUSSION

Although growth of smaller arterioles, venules, and capillaries in adult animals and later in development is largely driven by the needs of local tissues for oxygen and nutrients, and these vessels develop without reproducible patterns, the formation of major vessels during early development occurs highly stereotypically and reproducibly and is strongly guided by intrinsic genetic programs (Weinstein, 1999). We have shown in previous studies that major vascular networks that assemble during early development such as trunk intersegmental vessels (Isogai et al., 2001), the hindbrain central arteries (Fujita et al., 2011), and trunk lymphatic vessels (Cha et al., 2012; Jung et al., 2017) all form in a highly stereotyped, choreographed, and reproducible manner, and that the choreographed patterning of these vascular networks is guided by genetically programmed vascular guidance cues (Cha et al., 2012; Fujita et al., 2011; Lim et al., 2011; Torres-Vázquez et al., 2004). In this study we have shown that development of the major vessels of the pectoral fin vascular network occurs in a similarly choreographed and reproducible manner (**Fig. 9**), suggesting that assembly of these vessels is also likely guided by defined genetically programmed cues, although at this point these cues remain undiscovered.The first step in pectoral vessel formation is emergence of dorsal and ventral sprouts from the posterior cardinal vein. The timing and positioning of these two sprouts and their growth along the fin margin is quite reproducible, again suggesting directed guidance by defined cues. A positive cue from the apical ectodermal ridge (AER) along the outer margin of the developing fin would perhaps be sufficient to direct formation of two dorsally and ventrally positioned sprouts in the vicinity of the fin margin, perhaps in conjunction with additional negative cues from the central developing endoskeletal disk (ED; see **Fig. 3A-C**). As previously noted (Yano et al., 2012), the primitive pectoral artery (aka “circumferential blood vessel of the fin bud”) grows precisely along and defines the border between the proximal fin containing the endoskeletal disk (ED) and the distal apical fold (AF), a broad distal extension of the fish apical ectodermal ridge (Yano et al., 2012). Again, the positive and/or negative cues directing circumferential vessel growth along this corridor are not known, although there are many different secreted or membrane bound factors or localized matrices found along this corridor or in the adjacent endoskeletal disk and apical fold tissues (Mateus et al., 2020). Angiogenic cues such as VEGFA, FGF, and BMP are expressed in the developing fin bud and VEGF signaling has already been shown to be required for formation of the primitive pectoral artery (Habeck et al., 2002; Mateus et al., 2020; Nomura et al., 2006), but further studies would be needed to define which of these or other cues are required and how they work together to direct sprouting and circumferential outgrowth of the primitive pectoral artery. Given its precise positioning, it is also possible that the primitive pectoral artery itself plays a role in helping delimit and/or maintain the ED/EF boundary and/or in promoting outgrowth of the fin bud. This possibility could also be tested by limb-specific inhibition of vessel formation, although accomplishing this without the confounding problem of interfering with vessel development systemically would be extremely challenging.

**Figure 9.**
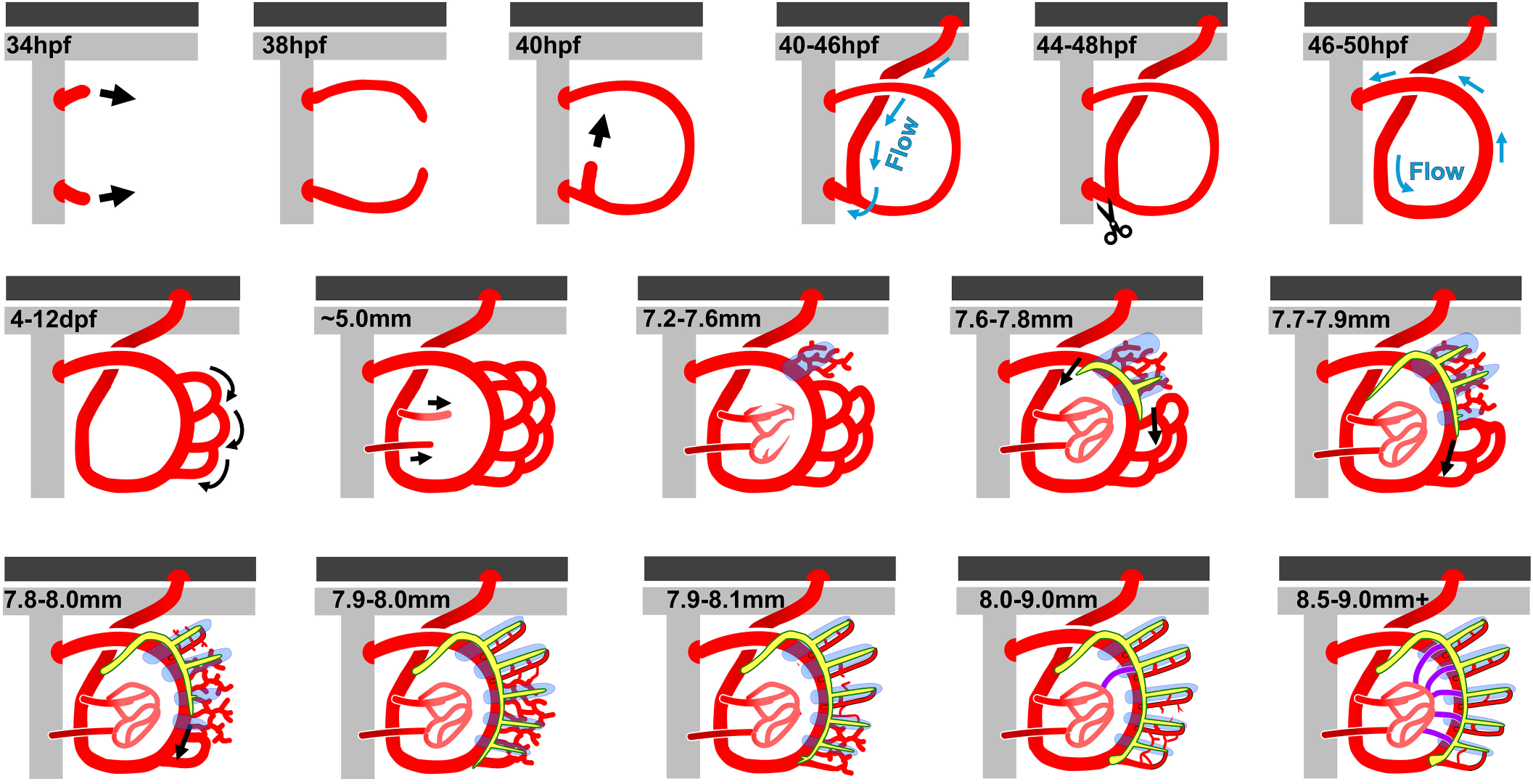
Schematic Diagram of Pectoral Fin Vascular Development. Schematic diagrams illustrating stages of pectoral fin vascular development from emergence of the first pectoral vessel sprouts from the CCV to formation of a complex vascular network that includes the major vascular pathways found in the adult pectoral fin. After the first week of development larval length is used instead of age to stage animals, as this more accurately correlates with developmental events (see **Fig. S4**)

Once formation of the circumferential loop of the primitive pectoral artery is completed, it becomes connected to arterial feed from the dorsal aorta by a fascinating process that involves extensive, long-distance angiogenic growth of a new vessel some 300 microns (**Fig. 4A-C**). This new vessel segment grows all the way from the ventral base of the developing fin bud via a highly reproducible path along the side of the fish and then between the base of the second and third somites before connecting to the dorsal aorta at the base of the second intersegmental artery (or in some cases actually attaching directly to the ventral-most part of the second intersegmental artery). Again, the local cues directing the growth of this vessel along its stereotyped and reproducible path remain to be determined. Several nerves track nearby along the lateral sides of the fish to the pectoral fin (Banerjee et al., 2015; Okamoto and Kuwada, 1991), but these nerves do not appear spatially closely linked to the growth of this vessel, suggesting they do not play a role in guiding its growth (data not shown). However, a substantial part of the path of this vessel does run in very close apposition to the anterior pronephric tubules. The developing pronephros has been shown to strongly express VEGF during stages of development just prior to and during growth of the vessel linking the pectoral vasculature to the aorta (Liang et al., 2001), suggesting the pronephros could be providing cues to direct at least part of the growth of this vessel. Although this idea could in theory be tested by specific ablation of the pronephros or pronephros-specific suppression of VEGF, the necessary transgenic tools for driving early expression in the appropriate regions of the anterior pronephros are not currently available in the fish.

Once the arterial connection is made, pectoral fin blood flow begins, but in most cases, blood initially drains directly into the common cardinal vein via a ventral shunt, bypassing the circumferential loop of the primitive pectoral artery which initially remains unperfused (**Fig. 4G**). However, within minutes to hours this shunt reproducibly disconnects, and all blood flow is re-routed through the remaining circumferential portions of the primitive pectoral vasculature (**Fig. 4H,I**). A large variety of studies, including in zebrafish and mouse, have shown that flow is a critical signal for vascular remodeling (Isogai et al., 2001; Kochhan et al., 2013; Lucitti et al., 2007) and historically vessel flow and resulting shear stress level have been thought to be an important modulator of vessel caliber and vessel retention, with high flow rates resulting in vessel enlargement/retention and low flow leading to vessel size decrease and/or regression (Baeyens and Schwartz, 2016; Kamiya and Togawa, 1980; Langille, 1996; Langille and O’Donnell, 1986; Tuttle et al., 2001), although it has been pointed out that unchecked vessel growth due to increased flow would be counterproductive and lead to non-functioning vascular networks (Pries et al., 2010), and recent studies have reported a variety of mechanisms limiting vessel growth and/or directing or restricting vessel connections, notably including the TGF-beta and Notch pathways (Ito et al., 2009; Phng et al., 2009; Suchting et al., 2007). In the developing pectoral fin a vessel connection with robust flow (the ventral shunt to the CCV) is disconnected and lost in order to send blood through a vessel that initially has no flow at all and that initially may not even be fully lumenized (i.e., the remaining portions of the circumferential pectoral artery), creating a longer (and presumably higher-resistance) path for blood flow than the shunt that it replaces. This suggests that ventral fin shunt disconnection and rerouting of blood flow through the remainder of the pectoral vasculature is actively driven by specific programmed disconnection cues or mechanisms, not as a “passive” result of flow dynamics. Although the nature of such disconnection cues or mechanisms remains unclear, Notch and TGF-beta ligand expression in the developing fin may suggest potential roles for these pathways.

Formation of a highly elaborated plexus at the distal tip of the fin begins at approximately one week post-fertilization (**Fig. 5**). As condensation of the fin ray bones initiates, this distal plexus then becomes progressively remodeled in a dorsal-to-ventral sequence in coordination with the dorsal-to-ventral development of each successive fin ray. This remodeling involves retention, growth, and enlargement of fin ray-associated vessels and regression of most non-fin ray-associated vessels (**Fig. 5M**). This series of events resembles the formation and subsequent chondrogenesis-associated remodeling of a distal vascular plexus in the developing human limb forelimb to form a simplified pattern of major vessels associated with the developing limb skeleton (Bates et al., 2002; Rodríguez-Niedenführ et al., 2001). In the fish there is minimal flow through much of this plexus during early remodeling stages, suggesting flow is not a significant driver of this process. Again, it seems likely that signals associated with developing bone or bone-associated cells are likely to be important proangiogenic cues for fin ray vascular development. During regeneration of the zebrafish fin chemokine signals associated with newly forming bone appear to be important for directing vascular cell migration and organization into new fin-ray associated vessels (Sivaraj and Adams, 2016; Thorimbert et al., 2015; Xu et al., 2014). It remains to be seen whether chemokine signaling plays a similar role in helping direct vessel remodeling during normal fin development and/or whether other cues are required. It also remains unclear whether, conversely, formation of the distal vascular plexus and subsequent fin ray vasculature play a role in chondro/osteogenesis and subsequent growth and development of the fin rays (Akiva et al., 2019). The roles of each of these tissues in the development of the other could be tested by specific disruption of fin-associated bone or vessel development, but the fin-specific manipulation of these tissues that would be needed to avoid the confounding effects of systemic disruption of these developing tissues is at present difficult or impossible to carry out in the fish. Remodeling of the distal fin vascular plexus to form the complex adult fin ray vasculature also involves specification of separated, parallel arterial and venous fin ray vascular networks (**Fig. 6**). A sprout initiating the dorsal part of the nascent arterial network appears to fairly reliably emerge from the vicinity of the first, most dorsal condensing pectoral fin ray bone, and kdrl-positive fin ray arteries then grow progressively along with and through the center of each developing fin ray. This suggests that the developing fin rays are providing cues not just for vessel growth but also for vessel “arterialization,” particularly in the central channel between the paired fin ray bones where fin ray arteries form. There are known features contained within the center channel of the fin ray such as fibroblasts that line the bone and the actinotrichia which may be providing these cues (Akiva et al., 2019; Huang et al., 2009). Chemokine signaling could again be providing an important cue. It is also interesting to note that VEGF itself promotes both vessel growth and, at higher levels, arterial differentiation (Weinstein and Lawson, 2002).

The processes and potential cues directing them described above are only a portion of those we have reported in this manuscript (**Fig. 9**), and our detailed description is probably itself only a partial glimpse at the full complex sequence of events needed to assemble the pectoral fin vascular network. In the future, understanding the molecular cues and mechanisms directing the fascinating stereotyped developmental processes required to for pectoral fin vascular development will require additional in-depth experimental studies employing cutting-edge tools and methods not yet available in the fish, such as tissue-specific optogenetic activation to specifically manipulate expression in the fin or in fin bone, endothelium, or other fin cell or tissue types. While they will be extremely challenging, successful completion of these sorts of future studies will not only provide important insights into mechanisms directing the assembly of vascular networks during development, but also potentially provide new therapeutic targets for interventions to treat cardiovascular disease, vascular malformations, and cancer.

## ACKNOWLEDGMENTS

We would like to thank Andrew Davis, Van Pham and Biniam Sileshi, as well as the rest of the current and former Weinstein Lab members who have given their invaluable time to help prepare this manuscript for publication. This work was supported by the intramural program of the *Eunice Kennedy Shriver* National Institute of Child Health and Human Development, National Institutes of Health (ZIA-HD001011 and ZIA-HD008915, to BMW).

## MATERIALS AND METHODS

### Fish Husbandry and Fish Strains

Fish were housed in a large zebrafish dedicated recirculating aquaculture facility (4 separate 22,000L systems) in 6L and 1.8L tanks. Fry were fed rotifers and adults were fed Gemma Micro 300 (Skretting) once per day. Water quality parameters were routinely measured, and appropriate measures were taken to maintain water quality stability (water quality data available upon request). The following transgenic fish lines were used for this study: *Tg(fli1a:egfp)^y1^* (Lawson and Weinstein, 2002), *Tg(mrc1a:egfp)^y251^* (Jung et al., 2017), *Tg(kdrl:mcherry)^y206^* (Gore et al., 2011), *Tg(kdrl:egfp)^s843^* (Jin et al., 2005), *Tg(gata1:dsred)^sd2Tg^* (Traver et al., 2003), and *Tg(Ola.Sp7:mCherryEco.NfsB)^pd46^* (aka *Tg(sp7:mcherry-ntr),* previously known as *Tg(osterix:mCherry-NTRo)^pd46^*, (Singh et al., 2012). Some of the lines imaged were maintained and imaged in a *casper* (*roy, nacre* double mutant) (White et al., 2008) genetic background in order to increase clarity for visualization of the pectoral fin due to melanocytes that develop on the fin itself as well as around the embryonic kidney.

### Image Acquisition

Confocal images were acquired primarily with a Nikon Ti2 inverted microscope with Yokogawa CSU-W1 spinning disk confocal, (Hamamatsu Orca Flash 4 v3 camera with the following Nikon objectives: 4X Air 0.2 N.A., 10X Air 0.45 N.A., 20X water immersion 0.95 N.A., and 25X silicone immersion 1.05 NA, 40X water immersion 1.15 NA, 60X water immersion 1.20 N.A.). A minority of images were taken using a Zeiss 880 with Fast Airyscan using a 40x Water Immersion or 10x air lens. Stereo microscope images were acquired using a Leica M205 microscope using MultiFocus focus stacking, or on a Leica M165 microscope with Leica DFC 7000T camera. For bone autofluorescence fish were illuminated with 405 nm laser excitation and imaged for 461 nm fluorescence emission (Castranova et al., 2021).

### Image Processing

Images were processed using Nikon Elements and Photoshop. Unless otherwise specified, maximum intensity projections of confocal stacks are shown. Focus stacking of confocal images with DIC was done using Nikon Elements EDF (Extended Depth of Focus). 3D rotation movies and time-lapse movies were made using Nikon Elements and exported to Adobe Premiere Pro CC 2019. Adobe Premiere Pro CC 2019 and Adobe Photoshop CC 2019 were used to add labels and arrows to movies and to add coloring or pseudo-coloring. Schematics were made using Adobe Photoshop CC 2019 and Adobe Illustrator CC 2019

### Fluorescence Quantification

Images quantified for relative fluorescence were generated using Imaris 9.1 by taking a flattened image of a max intensity 3D reconstruction of a z-stack. The flattened images were then quantified in Adobe Photoshop CC2019 by selecting relevant vascular beds in the indicated regions and measuring the average pixel intensity. To remove the influence of varying vascular density and region size, non-vascular pixels were excluded by threshold-including pixels with intensity numbers above background noise.

### Time Lapse Imaging and Shunt Quantification

Fish from 2 dpf to 4 dpf were imaged live by anesthetizing them in 168 mg/L tricaine in system water and mounting them in the common lateral position in 0.8% low melting point agarose in a 35mm glass bottomed petri dish (MatTek # P35-1.5-14-C). These fish were imaged for the indicated time interval for 36 hours on the Nikon W1 confocal as mentioned above. Shunting was quantified from live time-lapse imaging every 20 minutes of 12 *Tg(kdrl:egfp), Tg(gata1:dsred)* double transgenic animals imaged from ∼58 hpf (before flow onset) until 76 hpf. DsRed-positive red blood cells were monitored to determine whether they were shunting to the CCV or passing through the full primary arc of the primitive pectoral artery.

### Mounting and Imaging Later Larval and Adult Fish

To image larval zebrafish at stages from approximately 5 dpf to one month of age, fish were anesthetized in 168mg/L tricaine in E3 media and mounted laterally in 0.8% low melting point agarose in a 35 mm glass bottomed petri dish (MatTek # P35-1.5-14-C) for pectoral fin imaging. In some experiments animals were pre-anesthetized in 79mg/L tricaine in system water, where the fish were kept responsive to touch to await mounting, then transferred to 126 mg/L tricaine in system water for mounting and imaging as described above. Animals repeatedly imaged at multiple time points were gently recovered from the agarose and returned to fresh E3where they were gently continuously swirled until conscious and actively swimming and then returned to their tank on the system.

Fish older than one month were mounted for imaging by anesthetizing them in 126-168 mg/L tricaine in system water and then placed into a slit in a sponge (Jaece Identi-Plugs L800-D) moistened with tricaine water, cut into a rectangle to fit inside a single chamber imaging dish (Lab-TekII #155360). The sponge containing the fish was placed ‘belly-down” into the imaging chamber and the chamber was filled with tricaine water. Fish lengths are reported as standard length from the tip of the snout to most proximal part of the fork between caudal fin lobes.

Animals repeatedly imaged at multiple time points were recovered from the agarose after collecting images at each time point and returned to fresh E3.

