## Supplemental Information for "Anatomy and Development of the Pectoral Fin Vascular Network in the Zebrafish"

#### SUPPLEMENTAL FIGURE LEGENDS

##### Supplemental Figure 1. Sex-specific vascular features of the adult zebrafish pectoral fin.

**(A-D)** Tiled confocal images of vessels in pectoral fins dissected from *Tg(fli1a:egfp)* transgenic adult male (A,C) and female (B,D) zebrafish. Boxes in A and B indicate the approximate areas shown in the magnified images in panels C and D, respectively. Greater numbers of inter-ray vessels in are present in male pectoral fins. **(E)** Quantitation of the number of inter-ray vessels linking different sets of fin rays in adult male or female zebrafish. **(F)** Higher magnification confocal image of the same *Tg(kdrl:mcherry)* adult zebrafish fin as in **Fig. 2B**, showing kdrl:mcherry-positive arterial blood vessels (red, arrowheads) running between autofluorescent bones (blue) to interconnect the proximal and fin ray vascular networks. **(G-M)** Tiled confocal images of vessels in pectoral fins dissected from *Tg(osx:ntr-mcherry), Tg(kdrl:egfp)* double transgenic adult female (G-J) and male (K-M) zebrafish, with mCherry-positive bone in red and EGFP-positive vessels in green. The box on the image of an adult female fin in panel G indicates the approximate location of the magnified images shown in panels H-J. Panels K-M show magnified images from the comparable location in a separately imaged adult male fin, showing a complex network of vessels on the outside of the fin ray bone associated with the male breeding tubercles that are not present in the female fin. Scale bars = 1 mm, A; 200 µm , F-H; 500 µm, I; 30 µm, J-L; 40 µm, M-O.

##### Supplemental Figure 2. Quantitation of shunt formation and persistence

(A) A graph showing the total percentage of embryos with no flow, shunting flow, and normal flow, at the indicated time post fertilization. *n*=11. (B) Two images of the ventral arm of the primary pectoral fin artery in a 4 dpf *Tg(kdrl:egfp)* embryo with the ventral arterial sprout (VS) and ventral primary arc arm (VA) false colored in magenta and the common cardinal vein (CCV) in green. Left, lateral view. Right, ventral view with a blue arrowhead indicating the separation of the ventral arm from the CCV. Scale bars = 20 µm.

##### Supplemental Figure 3. Pectoral fin image series.

(**A-O**) Confocal images of pectoral fin distal plexus vascular development from 9 dpf to 26 dpf (4.3 mm to 10.5 mm) in a single representative *Tg(kdrl:egfp), Tg(mrc1a:egfp)* double transgenic zebrafish larva, with kdrl:mCherry in red and mrc1a:egfp in green. (**P,Q**) Confocal images of the pectoral fin distal vascular plexus in a 21 dpf *Tg(sp7:mcherry-ntr), Tg(kdrl:egfp)* double transgenic zebrafish, showing bone in magenta (P) and vessels in green (P,Q). White dashed lines in panel Q show the position of the bones, red and magenta dashed lines demarcate vascular areas in ray 1 and between rays 1 and 2, respectively. (**R**) Quantitation of kdrl:egfp fluorescence intensity of vessels in fin ray 1 and between fin rays 1 and 2. (**S-U**) Confocal images of the pectoral fin distal vascular plexus in 21 dpf *Tg(sp7:mcherry-ntr), Tg(fli1a:egfp)* double transgenic zebrafish, showing bone in red (S,T) and vessels in green (S-U). Panel S shows a lower-magnification overview of a distal vascular plexus, panels T and U show a higher magnification image of the tip of three fin rays from a separate fish. In panel U arrowheads indicate arteries, arrows indicate veins, and initial sprouts developing that will give rise to the dorsal paired fin ray veins are note with asterisks. All scale bars = 50 μm, except panel S, which is 100 μm.

**Supplemental Figure 4. Quantitation of the timing of developmental milestones in the pectoral fin vasculature.**

The two graphs on the left show the mean size measurement and chronological age of *Tg(kdrl:egfp), Tg(mrc1a:egfp)* double transgenic zebrafish larva imaged as the fish in **Supplemental Figure S3** and **Figure 5** as they achieved the indicated developmental milestones. All error bars indicate the standard deviation of the mean. *n* = 7 larva. The panels on the right show schematic representations of the events quantified in the graphs.

### Figure S1 - Paulissen et al.

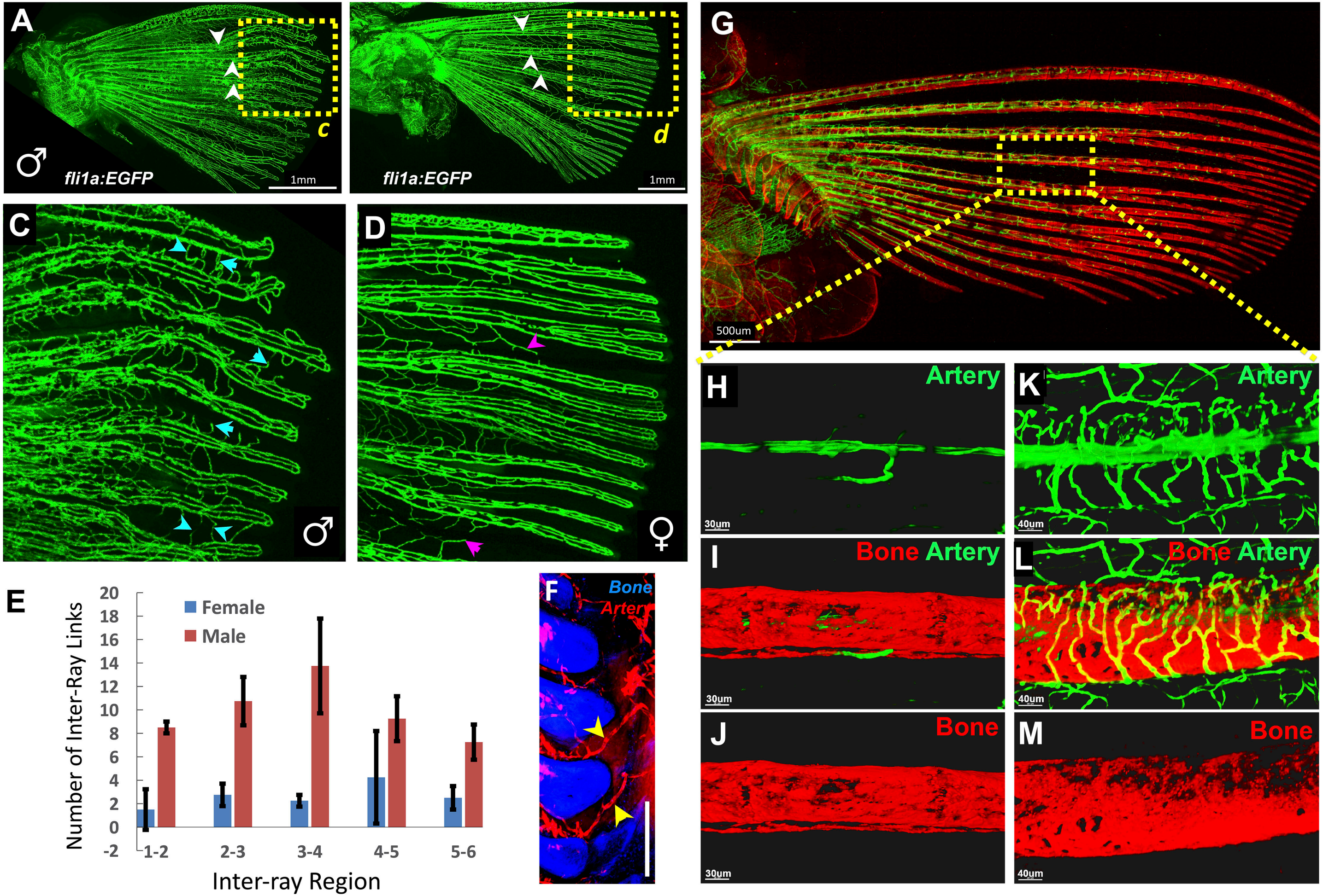

### Figure S2 - Paulissen et al.

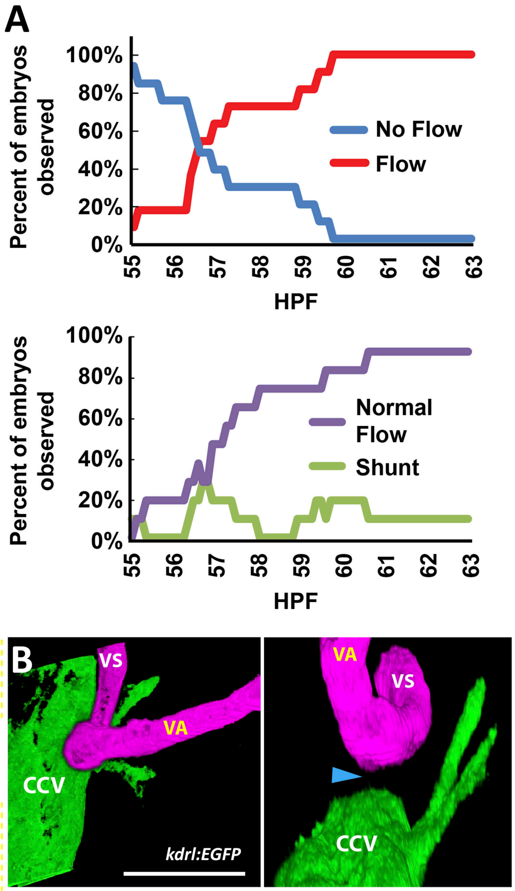

### Figure S3 - Paulissen et al.

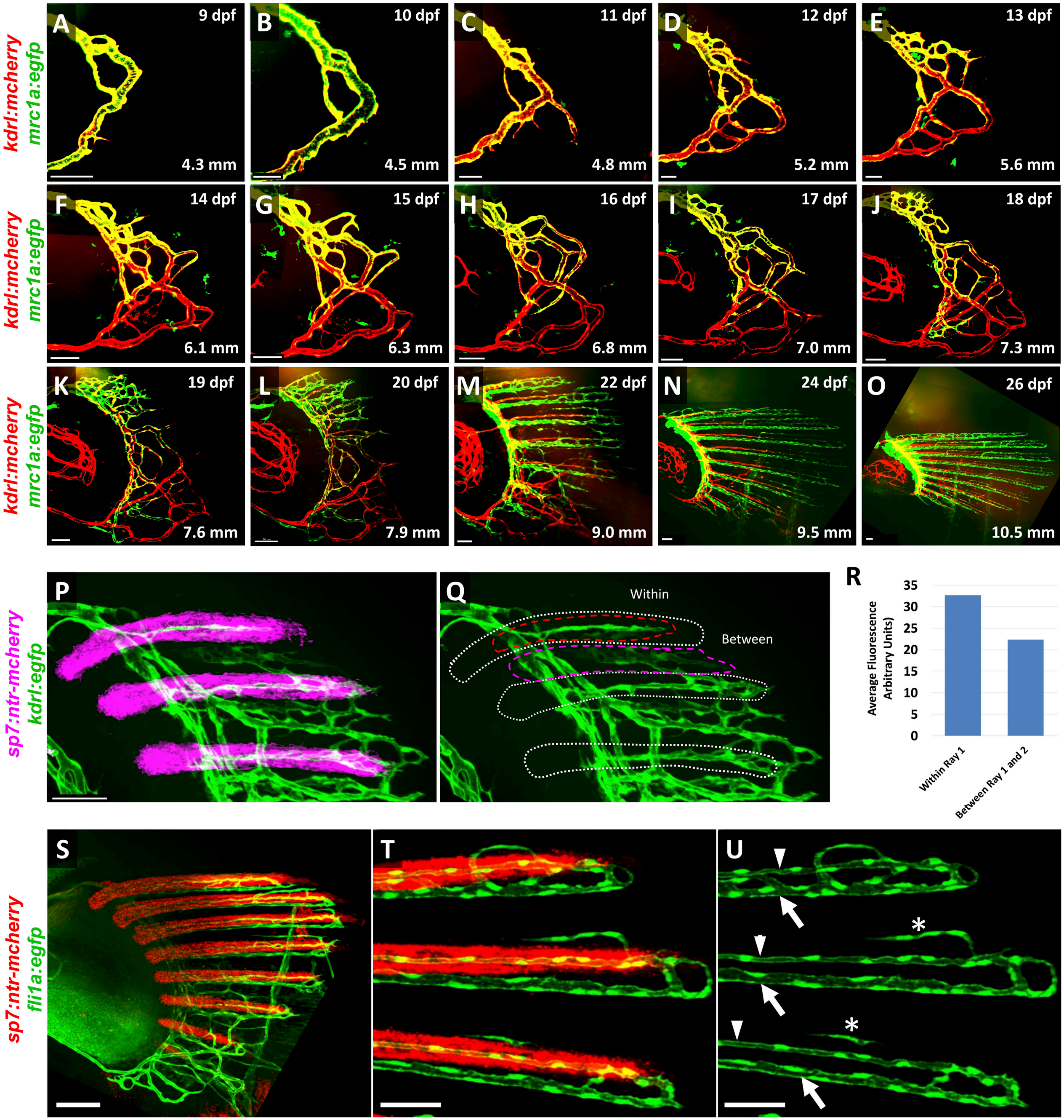

### Figure S4 - Paulissen et al.

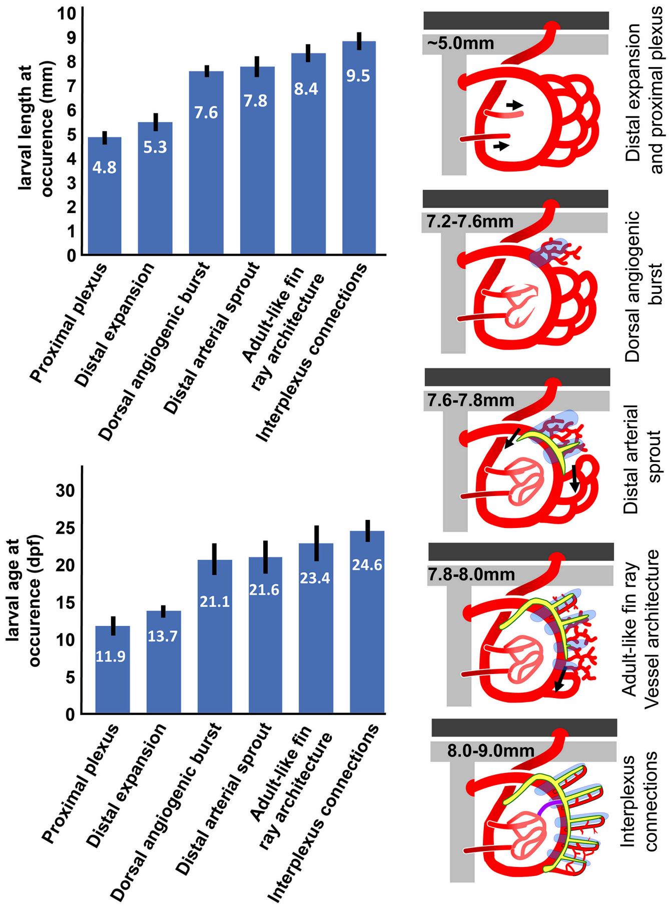

#### SUPPLEMENTAL MOVIES

**Supplemental Movie 1. Circulation in the adult zebrafish fin.**

High magnification transmitted light imaging of circulation through fin ray arteries and veins and inter-ray interlinking vessels of an adult zebrafish fin. Transmitted light, real time video.

**Supplemental Movie 2. Formation and arterial connection of the primitive pectoral artery.**

Time-lapse confocal imaging of a *Tg(kdrl:egfp)* transgenic zebrafish from 34-50 hpf, showing formation of the primary arc of the primitive pectoral artery and its connection to the dorsal aorta. The movie shows the complete sequence for the images shown in **Fig. 3D-F** and **Fig. 4D-F**.

**Supplemental Movie 3. Perfusion of the primitive pectoral artery.**

Time-lapse confocal imaging of a *Tg(kdrl:egfp)*, *Tg(gata1:dsred)* double transgenic zebrafish from 44-50 hpf, showing gata1:dsred marking active blood flow (left panel; pec fin flow colored red), kdrl:egfp showing both lumenized and unlumenized vessels (center panel; ventral connection to the DA colored green), and a merged gata:dsred (red)/kdrl:egfp (green) image (right panel). Flow initially shunts from the DA directly to the CCV, but then becomes re-routed through the full arc of the primitive pectoral artery. Each frame of the movie represents 20 minutes elapsed time. The movie shows the complete sequence for the images shown in **Fig. 4J-U**.

**Supplemental Movie 4. Initial shunting flow of blood flow from the DA to the CCV.**

3-D reconstructions of a confocal image stack of the fin region of a 2 day-old *Tg(kdrl:egfp)*, *Tg(gata1:dsred)* double transgenic zebrafish, showing flow initially shunting from the DA directly to the CCV with no perfusion of the main circumferential loop of the primitive pectoral artery. The movie shows the same fish as in **Movie 5** but at a slightly earlier time point.

**Supplemental Movie 5. Subsequent re-routing of blood flow through the primitive pectoral artery.**

3-D reconstructions of a confocal image stack of the fin region of the same 2 day-old *Tg(kdrl:egfp)*, *Tg(gata1:dsred)* double transgenic zebrafish as in **Movie 4**, showing that the shunt has severed and that flow from the DA has now been re-routed through the main circumferential loop of the primitive pectoral artery.

**Supplemental Movie 6. Distal vascular plexus of a 17 dpf larva**

3-D reconstructions of the same confocal image stack of the fin of a 17 day old (7.0 mm length) *Tg(kdrl:egfp)*, *Tg(gata1:dsred)* double transgenic zebrafish shown in **Fig. 5L**, with close-ups of the dorsal hypersprouting area of the distal vascular plexus.

**Supplemental Movie 7. Distal vascular plexus changes in an 18-22 dpf larva**

3-D reconstructions of the same confocal image stacks of the fin of an 18-22 day old (7.3-9.0 mm length *Tg(kdrl:egfp)*, *Tg(gata1:dsred)* double transgenic zebrafish as shown in **Fig. 5N-Q**, with close-ups of the dorsal hypersprouting area of the distal vascular plexus, including an addition 24 day old image of the same fish.

**Supplemental Movie 8. Development of the distal pectoral fin vasculature.**

Confocal time-series of the distal vascular plexus in the same *Tg(kdrl:mCherry), Tg(mrc1a:egfp)* double transgenic larva from day 9 through day 26 (4.3 mm to10.5 mm length). The movie shows the same images shown in **Fig. 5H-Q**.

**Supplemental Movie 9. Lateral and medial proximal plexuses of an 18 dpf zebrafish**

3D reconstructions of a confocal image stack of the fin of an 18 dpf (7.9 mm) *Tg(kdrl:mCherry), Tg(mrc1a:egfp)* double transgenic larva. The entire fin is initially shown, then only kdrl:mCherry positive arteries (red), then only the proximal plexus region. 90-degree rotations of the proximal plexus show that it actually consists of two parallel plexuses, a lateral plexus (red) and a medial plexus (false-colored blue).
